# TSC1 phosphorylation by lysosomal mTORC1 establishes a minimal autoregulatory feedback loop

**DOI:** 10.64898/2026.01.15.699678

**Authors:** Andreas Lamprakis, Constantinos Demetriades

**Author notes:** Correspondence (C.D.).

## Abstract

mTORC1 lies at the center of an intricate signaling network that allows cells to homeostatically respond to a multitude of intra- and extracellular cues. Although this network is dynamically rewired to prevent excessive mTORC1 activation under permissive conditions, the mechanisms that fine-tune its activity remain incompletely understood. Here, we identify TSC1 (tuberous sclerosis complex 1), a core subunit of the mTORC1-inhibitory TSC complex, as a direct mTORC1 substrate, thus establishing an autoregulatory circuit in mTOR signaling. Notably, TSC1 combines features of both canonical and non-canonical/lysosomal mTORC1 substrates: while its phosphorylation depends on Rheb activation and growth factor signaling, it also requires an intact lysosomal LAMTOR–Rag GTPase supercomplex. In turn, TSC1 phosphorylation selectively influences lysosomal mTORC1 signaling, as expression of a phospho-dead TSC1 mutant is associated with lower levels of TFEB phosphorylation, increased nuclear TFE3 translocation, and enhanced lysosome biogenesis, while phosphorylation of S6K1, a non-lysosomal canonical substrate, remains largely unaffected. Mechanistically, phosphorylation of TSC1 by mTORC1 promotes its stability, with a non-phosphorylatable mutant undergoing proteasomal degradation. In sum, these findings reveal a minimal feedback loop within the mTOR network that orchestrates compartmentalized signaling to selectively control the activation of processes downstream of lysosomal mTORC1 signaling.

## Introduction

The mammalian/mechanistic target of rapamycin (mTORC1) is a protein complex with kinase activity that integrates a wide variety of environmental cues such as growth factors, energy levels, oxygen status, and amino acids, to regulate cellular growth. mTORC1 regulates multiple diverse downstream processes, including protein synthesis, ribosome biogenesis, autophagy, and lysosome biogenesis, by phosphorylating its downstream substrates ^1–3^. mTORC1 largely assimilates these diverse stimuli via signaling pathways that relay the status of these conditions through specific phosphorylation events on the tuberous sclerosis complex (TSC) ^4^.

The heterotrimeric TSC complex is the major negative regulator of mTORC1 activity and comprises the proteins TSC1, TSC2 (also known as Hamartin and Tuberin, respectively), and the auxiliary component TBC1D7 (TBC1 domain family member 7) in a 2:2:1 stoichiometry ^5,6^. Mutations in TSC1 or TSC2 are linked to tuberous sclerosis complex (TSC), a rare genetic disorder characterized by the formation of benign tumors (hamartomas) in multiple organs, including the brain, skin, kidneys, and heart, as well as neurological symptoms such as epilepsy and developmental delay ^7,8^. The TSC2 subunit of the complex contains a C-terminal GTPase-activating protein (GAP) domain that is responsible for GTP hydrolysis of the RHEB GTPase (Ras homolog enriched in brain), the direct upstream activator of mTORC1 ^9,10^. While TSC2 possesses GAP activity, TSC1 is an obligate regulatory partner that promotes TSC complex activity by stabilizing TSC2 ^11,12^. More recently, TSC1 was shown to be a determining factor in the lysosomal localization of the complex ^13^. Lysosomal recruitment of the TSC complex is integral to its ability to suppress mTORC1 signaling. Under conditions of nutrient or growth factor deprivation, as well as in response to various cellular stresses, the TSC complex translocates to the lysosomal surface, where it encounters RHEB and converts it from the active GTP-bound form to the inactive GDP-bound form, thereby inhibiting mTORC1 activity ^9,14–17^. Notably, genetic deletion of either TSC1 or TSC2 leads to constitutive activation of mTORC1, making it unresponsive to perturbations in cellular growth conditions, thus highlighting the crucial role of the TSC complex in orchestrating mTORC1 activity ^7,8,18^.

Given the vast array of biological processes controlled by mTORC1, fine-tuning its activity is of utmost importance for cellular physiology. To ensure that the pathway is kept under homeostatic control, feedback loops exist to maintain optimal signaling output of mTORC1 ^19^. In several cases, feedback regulation in the pathway is mediated via the deposition of phosphorylation marks on the TSC complex by distal signaling kinases ^20^. For instance, when mTORC1 is suppressed, feedback regulation of upstream signaling effectors, involving lipid second messengers and mTORC2, leads to compensatory activation of the PI3K-AKT pathway upstream of the TSC complex to ultimately restore mTORC1 basal activity ^21–24^. Conversely, when mTORC1 is constitutively active, the exact feedback mechanisms are engaged to keep the pathway homeostatically balanced by terminating upstream signaling events. Although many of the distal signaling networks that are involved in the feedback regulation of the mTORC1 pathway upon its hyperactivation have been described before, the effect on more proximal components of the mTOR pathway is thus far unexplored.

Here, by using genetic, pharmacological, and nutritional means to perturb mTORC1 activity, we identify TSC1 as a novel mTORC1 substrate. TSC1 phosphorylation depends on an intact lysosomal nutrient-sensing machinery and responds to changes in amino acid, glucose, and growth factor availability. Lack of mTORC1-dependent phosphorylation on TSC1 decreases its stability and preferentially affects lysosomal mTORC1 activity, while the phosphorylation of S6K1, a non-lysosomal mTORC1 target, remains unchanged. Overall, our results reveal an autoregulatory feedback loop, shaped by the TSC, RHEB, and mTORC1 signaling modules, that operates to fine-tune substrate-specific mTORC1 signaling in cells.

## Results

### mTORC1 activation drives TSC1 phosphorylation

RHEB is the immediate upstream positive regulator of mTORC1, and RHEB overexpression systems are commonly used to assess the consequences of mTORC1 activation ^25^. Using such models, we previously examined the role of RHEB in the regulation of mTORC1 by nutrients and osmostress ^17,25^. In the course of these studies, we noticed that transient expression of wild-type RHEB or of the constitutively active RHEB S16H (RHEB^S16H^) mutant ^26^ in HEK293FT cells led to a pronounced mobility shift of TSC1 on SDS-PAGE, coinciding with elevated mTORC1 activity, as evidenced by an increase in the phosphorylation of its canonical substrate S6K1 (Figure 1A). This effect was absent in cells expressing an inactive RHEB mutant (I39K), indicating that the perceived upshift in TSC1 migration was the result of increased RHEB activity rather than its elevated protein levels. Similar results were obtained in cells expressing RHEBL1, a functionally redundant paralog of RHEB, as well as an oncogenic RHEB mutant (Y35N) (Figure S1A-B) ^27,28^.

**Figure 1.**
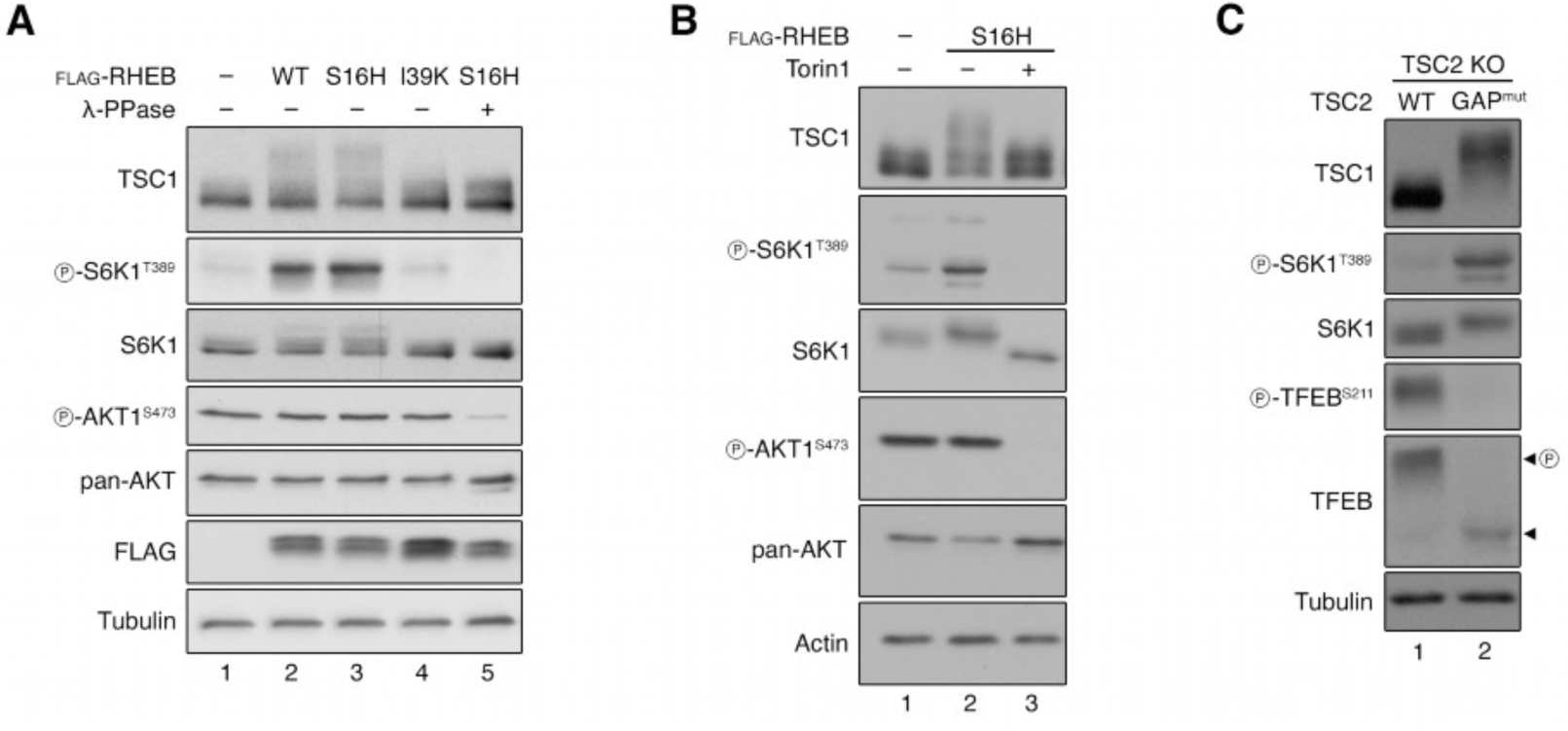
mTORC1 hyperactivation drives TSC1 phosphorylation. **(A)** Immunoblots with lysates from HEK293FT cells transiently expressing FLAG-tagged wild-type RHEB (WT), active RHEB^S16H^ (S16H), or inactive RHEB^I39K^ (I39K), or an empty vector as control (–), probed with the indicated antibodies. Lambda phosphatase (λ-PPase) treatment was used to remove phosphorylation from the indicated proteins. n = 3 independent experiments. **(B)** Immunoblots with lysates from HEK293FT cells transiently expressing FLAG-tagged RHEB^S16H^ or an empty vector, treated with Torin1 (250 nM, 1 h) or DMSO as control, probed with the indicated antibodies. n = 3 independent experiments. **(C)** Immunoblots with lysates from TSC2 KO HEK293FT cells stably expressing wild-type human TSC2 (WT) or the N1643K GAP-inactive hTSC2 mutant (GAP^mut^), probed with the indicated antibodies. n = 3 independent experiments. See also Figures S1 and S2.

Phosphorylation-dependent electrophoretic mobility shift in SDS-PAGE is a common phenomenon in cell signaling studies. Indeed, treatment of cell lysates with λ-phosphatase abolished the TSC1 mobility shift, confirming that the RHEB-induced TSC1 modification is phosphorylation-dependent (Figure 1A). Furthermore, treatment of RHEB^S16H^-expressing cells with Torin1, a catalytic inhibitor of mTOR (Figure 1B), fully and rapidly restored TSC1 electrophoretic mobility within one hour, indicating that mTOR mediates the RHEB-dependent phosphorylation of TSC1. This was not due to the involvement of mTORC2, as Rheb^S16H^ overexpression—which promotes TSC1 hyperphosphorylation—did not cause detectable changes in AKT1 phosphorylation, a typical read-out of mTORC2 activity (Figures 1A-B).

To assess whether this effect is applicable also to different models of mTORC1 hyperactivation, we examined the mobility shift of TSC1 in TSC2 knockout (KO) cells. Surprisingly, despite strong mTORC1 hyperactivation in these cells, TSC1 phosphorylation remained unchanged, even when overexpressing RHEB^S16H^ (Figure S1C). Therefore, we hypothesized that the integrity of the TSC complex may be required for mTORC1-dependent phosphorylation of TSC1. Indeed, reconstitution of TSC2 KO cells with a GAP-inactive TSC2 mutant—in which endogenous RHEB and mTORC1 signaling is hyperactive, despite the presence of an intact TSC complex—led to strongly elevated TSC1 phosphorylation (Figure 1C). The upshift in TSC1 mobility was observed also in human osteosarcoma U2OS cancer cells exogenously expressing RHEB^S16H^, and in mouse embryonic fibroblasts (MEFs) stably expressing GAP-mutant TSC2, showing that this effect is not cell-type- or species-specific (Figures S2A-B).

### TSC1 is a direct, physiological mTORC1 substrate

To identify the mTORC1-dependent phosphorylation sites on TSC1, we combined endogenous TSC1 immunoprecipitation with mass spectrometry, using HEK293FT cells expressing RHEB^S16H^, treated with Torin1 of DMSO as control. These experiments led to the identification of two TSC1 peptides containing residues T1047 and S1080, respectively, whose phosphorylation was detected in RHEB^S16H^-expressing cells and reduced upon mTOR inhibition (Figure 2A and Table S1), suggesting that TSC1^T1047^ and TSC1^S1080^ may be phosphorylated by mTORC1.

**Figure 2.**
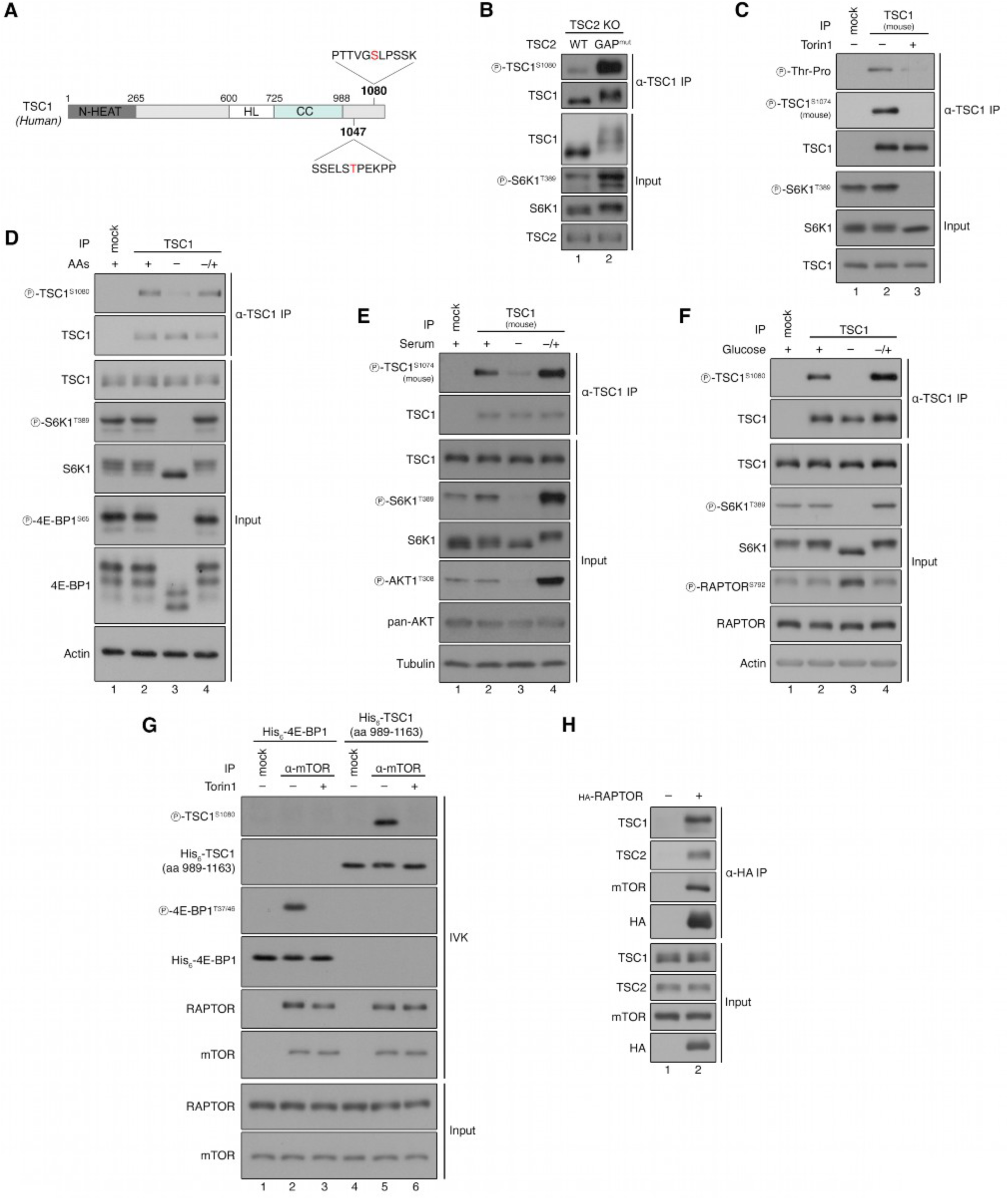
TSC1 is a direct, physiological mTORC1 substrate. **(A)** Schematic representation of the mTOR-dependent phosphorylation sites on human TSC1 protein, identified by phospho-proteomics. The amino acid sequence surrounding positions T1047 and S1080 is shown, as well as the position of the N-terminal HEAT (N-HEAT), the helical linker (HL) and the coiled-coil (CC) TSC1 domains. **(B)** Endogenous TSC1 immunoprecipitation (IP) from TSC2 KO HEK293FT cells stably expressing wild-type human TSC2 (WT) or the N1643K GAP-inactive hTSC2 mutant (GAP^mut^), followed by immunoblotting with the indicated antibodies. n = 3 independent experiments. **(C)** Endogenous TSC1 immunoprecipitation from MEFs treated with Torin1 (250 nM, 1 hr) or DMSO as control, followed by immunoblotting with the indicated antibodies. n = 3 independent experiments. **(D)** Endogenous TSC1 immunoprecipitation (IP) from HEK293FT cells treated with media containing or lacking amino acids (AAs), in basal (+), starvation (–), or add-back (–/+) conditions, followed by immunoblotting with the indicated antibodies. n = 3 independent experiments. **(E)** Endogenous TSC1 immunoprecipitation (IP) from MEFs treated with media containing or lacking serum, in basal (+), starvation (–), or add-back (–/+) conditions, followed by immunoblotting with the indicated antibodies. Note that S1080 in human TSC1 corresponds to S1074 in mouse TSC1. n = 3 independent experiments. **(F)** Endogenous TSC1 immunoprecipitation (IP) from HEK293FT cells treated with media containing or lacking glucose, in basal (+), starvation (–), or add-back (–/+) conditions, followed by immunoblotting with the indicated antibodies. n = 3 independent experiments. **(G)** *In vitro* kinase assays with endogenous mTOR immunopurified from HEK293FT cells, using recombinant His6-tagged TSC1^989-1163^ as a substrate. Recombinant His_6_-tagged 4E-BP1 was used as a positive control. Substrate phosphorylation detected by immunoblotting. Addition of Torin1 (250 nM) to the tubes 10 min before the initiation of the IVK reactions was used to confirm mTOR-dependent substrate phosphorylation. n = 2 independent experiments. **(H)** Co-immunoprecipitation experiments with lysates from HEK293FT cells transiently expressing HA-tagged RAPTOR or a control vector reveal binding of mTORC1 to endogenous TSC1 and TSC2. The input and anti-HA IP samples were analyzed by immunoblotting with the indicated antibodies. n = 3 independent experiments. See also Figures S3 and S4.

To look more closely into TSC1 phosphorylation by mTORC1, we next generated a phospho-specific antibody recognizing the epitope surrounding phospho-S1080 in human TSC1 (S1074 in mouse TSC1). Using TSC1 KO cells re-expressing wild-type TSC1 or a phospho-dead S1080A TSC1 mutant, we confirmed the specific detection of S1080 phosphorylation on immunopurified endogenous TSC1 (Figure S3). Moreover, an increase in TSC1^S1080^ phosphorylation correlated with the TSC1 upshift and elevated mTORC1 signaling in TSC2 KO cells expressing a TSC2 GAP-inactive mutant, consistent with our mass spectrometry data (Figure 2B). Further supporting that TSC1^T1047^ may be an mTORC1-dependent phosphosite, the sequence immediately downstream of T1047 is occupied by a proline residue, in line with mTOR being a proline-directed kinase (Figure 2A) ^29^. Indeed, using a phospho-threonine-proline antibody (that detects phosphorylated threonine residues only when followed by proline), we confirmed the mTOR-dependent phosphorylation on immunopurified TSC1 (Figure 2C).

A remarkable feature of the mTOR pathway is the wide range of intracellular and extracellular cues it integrates. To investigate whether physiological stresses that are known to control mTORC1 activity can also impinge on TSC1 phosphorylation, we starved cells from, or resupplemented them with amino acids (AAs), growth factors, or glucose. Resembling the behavior of well-described canonical mTORC1 substrates, like S6K1 and 4E-BP1, withdrawal of amino acids, growth factors, or glucose resulted in a robust decrease in TSC1 phosphorylation, followed by recovery upon resupplementation (Figures 2D-F).

We then investigated whether TSC1 may serve as a direct substrate of mTORC1. To test this, we performed *in vitro* kinase assays using immunopurified endogenous mTOR and a bacterially-expressed, C-terminal fragment of TSC1 (aa 989-1163), or recombinant 4E-BP1 as a positive control. These experiments showed that mTOR complexes can phosphorylate TSC1^S1080^ directly and specifically, as this modification was abolished entirely by Torin1 treatment (Figure 2G). Moreover, co-immunoprecipitation (co-IP) experiments revealed interactions between ectopically expressed HA-tagged RAPTOR, a unique accessory component of mTORC1 that is known to mediate canonical substrate recruitment, and endogenous TSC1 and TSC2 proteins (Figure 2H). Based on the premise that the presence of TSC2 is necessary for TSC1 phosphorylation by mTORC1 (Figure S1C), we reasoned that TSC2 may be the scaffold for the interaction between mTORC1 and the TSC complex. However, HA-tagged RAPTOR could readily interact with TSC1 also in TSC2 KO cells, or in cells expressing an N-terminally truncated TSC2 form that lacks the first 424 amino acids, encompassing its TSC1-binding domain (ΔT1BD) ^30^ (Figure S4). Thus, while TSC2 is essential for mTORC1 to phosphorylate TSC1, the TSC1-mTORC1 interaction can also occur independently of TSC2. Collectively, our data support a model wherein mTORC1 directly interacts with and phosphorylates TSC1, thereby establishing the latter as a previously unrecognized mTORC1 substrate.

### TSC1 phosphorylation requires intact lysosomal mTORC1 signaling

Under nutrient-replete conditions, the Rag GTPases (hereafter referred to as the Rags) promote the recruitment and activation of a fraction of mTORC1 on the lysosomal surface, thereby coupling amino acid availability to mTORC1 signaling ^31–34^. The Rags function as obligate heterodimers, consisting of either RagA or RagB bound to RagC or RagD, and the four possible Rag combinations have been shown to be functionally non-redundant in regulating mTORC1 activity toward its various substrates ^35^. In addition to their role in mTORC1 tethering, the Rag GTPases also mediate localization of the TSC complex to lysosomes upon amino acid withdrawal, primarily via interactions between RagA and TSC2 ^17^. Finally, recent studies revealed the spatial separation of mTORC1 signaling in cells, with lysosomal and non-lysosomal mTORC1 phosphorylating distinct substrates at each subcellular location ^33^.

To test whether lysosomal mTORC1 signaling is required for TSC1 phosphorylation, we transiently knocked down expression of the RagA/RagC or the RagB/RagD dimers in cells expressing GAP-mutant TSC2 that demonstrate mTORC1 hyperactivation and elevated TSC1 phosphorylation. Notably, knockdown of RagA/C—but not of RagB/D—could fully reverse the TSC1 upshift in these cells (Figure 3A), which is consistent with the previously-described role of RagA-containing dimers in mediating the lysosomal recruitment of the TSC ^17^. In contrast, the phosphorylation of S6K1, a cytoplasmic mTORC1 substrate, was largely unaffected by either Rag knockdown combination (Figure 3A), in agreement with recent work demonstrating the Rag- and lysosome-independent regulation of S6K1 phosphorylation downstream of mTORC1 ^33^.

**Figure 3.**
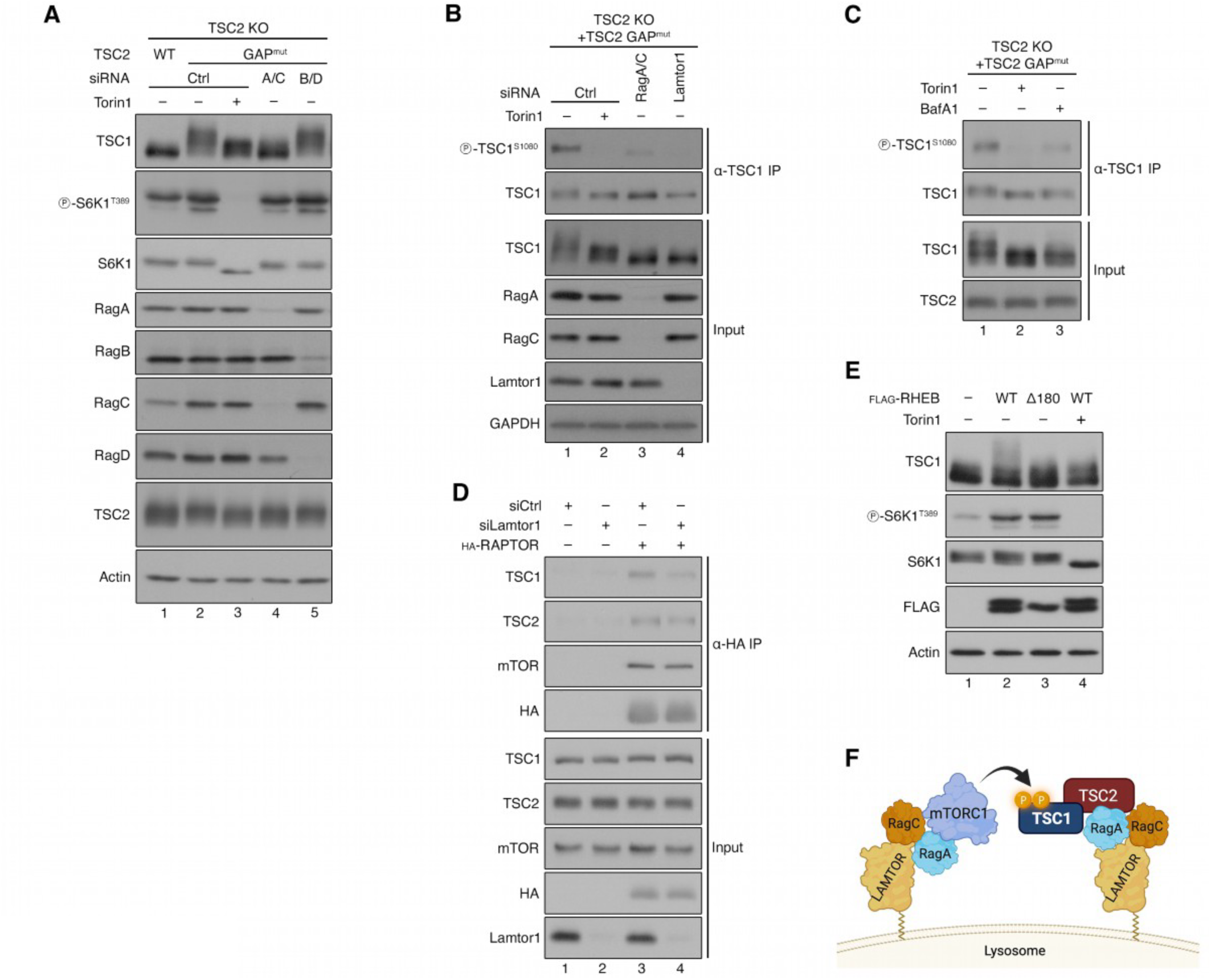
TSC1 phosphorylation requires intact lysosomal mTORC1 signaling. **(A)** Immunoblots with lysates from TSC2 KO HEK293FT cells stably expressing wild-type human TSC2 (WT) or the N1643K GAP-inactive hTSC2 mutant (GAP^mut^) and transfected with siRNAs targeting RagA/C, RagB/D, or Luciferase as a control, probed with the indicated antibodies. Torin1 (250 nM, 1 h) was used to inhibit mTOR. n = 3 independent experiments. **(B)** Endogenous TSC1 immunoprecipitation from TSC2 KO HEK293FT cells stably expressing the N1643K GAP-inactive hTSC2 mutant (GAP^mut^) and transfected with siRNAs targeting RagA/C, Lamtor1, or Luciferase as a control, probed with the indicated antibodies. Torin1 (250 nM, 1 h) was used to inhibit mTOR. n = 3 independent experiments. **(C)** Endogenous TSC1 immunoprecipitation from TSC2 KO HEK293FT cells stably expressing the N1643K GAP-inactive hTSC2 mutant (GAP^mut^), treated with Torin1 (250 nM, 1 h) or Bafilomycin A1 (100 nM, 8 h), probed with the indicated antibodies. n = 3 independent experiments. **(D)** Co-immunoprecipitation experiments with lysates from HEK293FT cells transiently expressing HA-tagged RAPTOR or a control vector and transfected with siRNAs targeting Lamtor1, or Luciferase as a control. The input and anti-HA IP samples were analyzed by immunoblotting with the indicated antibodies. n = 3 independent experiments. **(E)** Immunoblots with lysates from HEK293FT transiently expressing FLAG-tagged wild-type RHEB (WT) or the farnesylation-deficient RHEB^Δ180^ mutant (Δ180), treated with Torin1 (250 nM, 1 h) or DMSO as control, probed with the indicated antibodies. n = 2 independent experiments. **(F)** Schematic model illustrating the LAMTOR- and Rag-dependent phosphorylation of TSC1 on the lysosomal surface. See text for details.

As RagC-containing dimers dynamically cycle between the cytoplasm and the lysosomal surface in a nutrient-dependent manner ^35,36^, we next asked if TSC1 phosphorylation is a lysosomal event or occurs in a Rag-dependent manner elsewhere in the cell. To test this, we disrupted lysosomal anchoring of Rag GTPases by depleting Lamtor1, a core component of the LAMTOR (late endosomal/lysosomal adaptor, MAPK and mTOR activator) complex that scaffolds the Rags to the lysosomal membrane ^37–39^. RNAi-mediated silencing of Lamtor1 led to a strong reduction in TSC1^S1080^ phosphorylation, to a level comparable to Torin1 treatment (Figure 3B). Consistent results were obtained in cells treated with Bafilomycin A1 (BafA1), a macrolide antibiotic and potent v-ATPase inhibitor that perturbs lysosomal function and is known to specifically inhibit mTORC1 activity toward its lysosomal substrates, such as TFEB and TFE3, even in the presence of exogenous AAs ^33,40,41^ (Figure 3C). Finally, Lamtor1 knockdown in cells ectopically expressing HA-tagged RAPTOR weakened its association with the TSC (Figure 3D), indicating that the LAMTOR complex may be bridging mTORC1 and TSC1 for the latter to be phosphorylated on lysosomes.

Farnesylation of RHEB facilitates its anchorage to various endomembranes, including lysosomes ^42^. Interestingly, exogenous expression of a RHEB truncate lacking the C-terminal farnesylation motif (Δ180) was not able to drive TSC1 phosphorylation, while it potently enhanced the phosphorylation of S6K1, a cytoplasmic mTORC1 substrate (Figure 3E). Taken together, these findings demonstrate that TSC1 phosphorylation by mTORC1 requires an intact Rag-LAMTOR supercomplex, membrane-localized RHEB, and functional lysosomal mTORC1 signaling, thus highlighting TSC1 as a *bona fide* lysosomal mTORC1 substrate (Figure 3F).

### mTORC1-dependent phosphorylation stabilizes TSC1 and selectively regulates lysosomal mTORC1 signaling

To elucidate the impact of TSC1 phosphorylation, predicated on our knowledge of the phospho-acceptor residues, we generated TSC1 KO cells in which we transiently expressed either wild-type TSC1 or a non-phosphorylatable mutant harboring alanine substitutions for the respective mTORC1 sites T1047 and S1080 (hereafter TSC1^2A^) (Figure 2A). Strikingly, TSC1 protein levels were markedly reduced in cells expressing the phospho-deficient TSC1^2A^ mutant, suggesting that TSC1 phosphorylation promotes its stability. Treatment with the proteasome inhibitor MG132 restored TSC1^2A^ protein levels, indicating that mTORC1-dependent phosphorylation regulates TSC1 protein levels in a proteasomal-dependent manner (Figure 4A). Because TSC complex integrity contributes to the mutual stabilization of TSC1 and TSC2 ^5^, we speculated that the decrease in TSC1^2A^ protein levels may be the result of impaired interaction between the mutant TSC1 with TSC2. However, TSC1^WT^ or TSC1^2A^ showed similar binding to TSC2 in co-IP experiments, indicating that the phosphorylation-dependent effects on TSC1 protein stability do not involve changes in TSC complex integrity (Figure 4B). As an independent confirmation of the mTORC1-driven control of TSC1 protein stability, Torin1 treatment accelerated TSC1 turnover in cycloheximide (CHX)-treated cells (Figures 4C-D).

**Figure 4.**
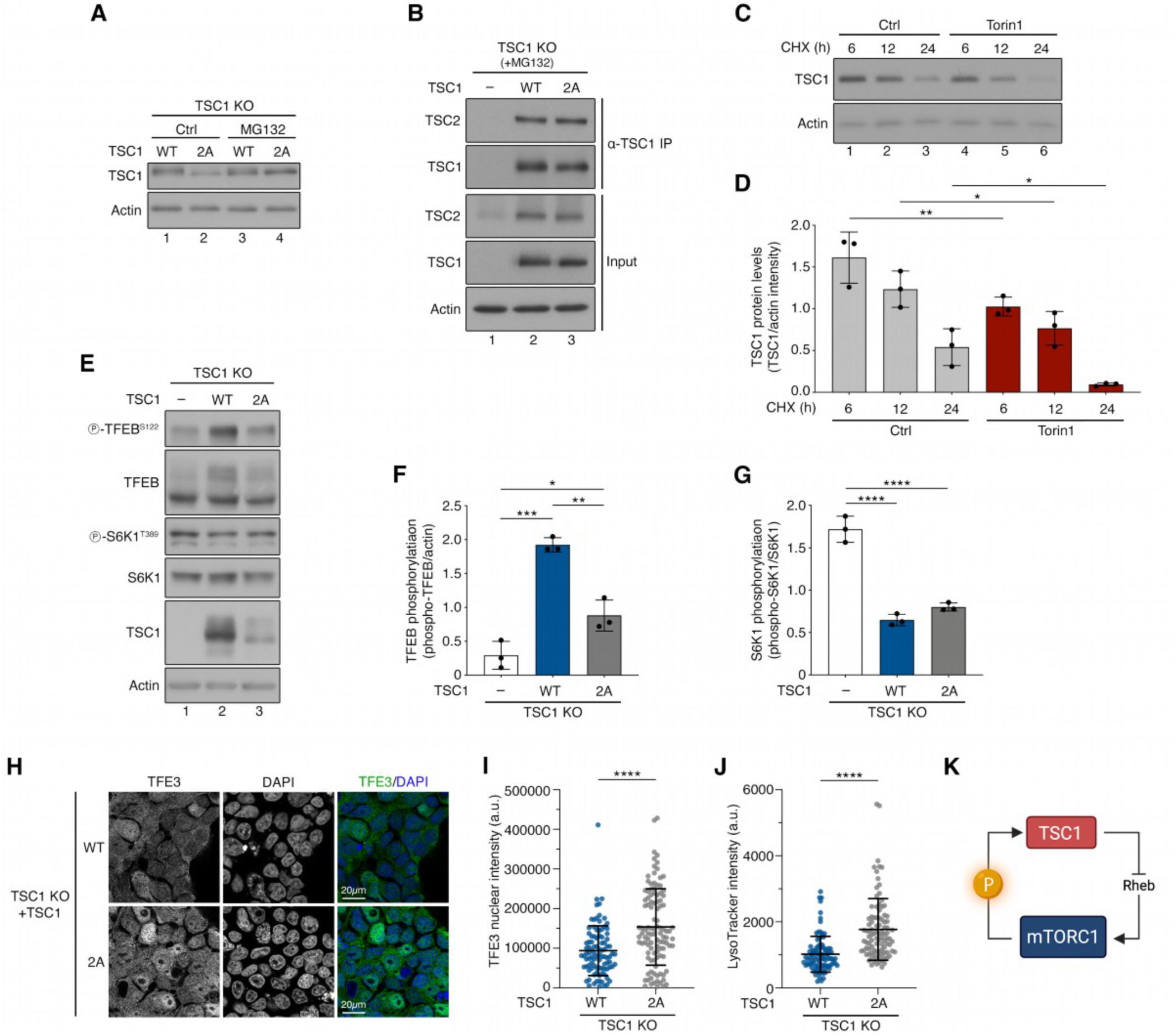
mTORC1-dependent TSC1 phosphorylation promotes its stability and modulates lysosomal signaling. **(A)** Immunoblots with lysates from TSC1 KO HEK293FT cells transiently expressing wild-type TSC1 (WT) or the phospho-dead TSC1^2A^ mutant (2A), treated with MG132 (10 μM, 8 h) or DMSO as control, probed with the indicated antibodies. n = 3 independent experiments. **(B)** TSC1 immunoprecipitation from TSC1 KO HEK293FT cells transiently expressing wild-type TSC1 (WT) or the phospho-dead TSC1^2A^ mutant (2A), followed by immunoblotting with the indicated antibodies. Cells were treated with MG132 (10 μM, 8 h) to block proteasomal degradation. n = 3 independent experiments. **(C-D)** Immunoblots with lysates from MEFs treated with cycloheximide (CHX, 100 μM) alone or in combination with Torin1 (250 nM) for the indicated time points (C). Quantification of TSC1 protein levels in (D). n = 3 independent experiments **(E-G)** Immunoblots with lysates from TSC1 KO HEK293FT cells transiently expressing wild-type TSC1 (WT) or the phospho-dead TSC1^2A^ mutant (2A), probed with the indicated antibodies (E). Quantification of TFEB phosphorylation and S6K1 phosphorylation in (F) and (G), respectively. n = 3 independent experiments. **(H-I)** TFE3 localization analysis in TSC1 KO HEK293FT cells transiently expressing wild-type TSC1 (WT) or the phospho-dead TSC1^2A^ mutant (2A), using confocal microscopy (H). Quantification of TFE3 nuclear intensity (arbitrary units, a.u.) in (I). n = 95-108 individual nuclei. **(J)** Quantification of LysoTracker signal intensity (arbitrary units, a.u.) from TSC1 KO HEK293FT cells transiently expressing wild-type TSC1 (WT) or the phospho-dead TSC1^2A^ mutant (2A). n = 96 individual cells. **(K)** Schematic model of the TSC1-mTORC1-TSC1 negative feedback loop. Elevated lysosomal mTORC1 activity drives TSC1 phosphorylation in specific residues that prevents its proteasome-mediated degradation. In turn, phosphorylated TSC1 preferentially downregulates lysosomal mTORC1 signaling, thus activating TFEB-dependent lysosome biogenesis (see also text for details). Data in graphs shown as mean ± SD. \**p* < 0.05, \*\**p* < 0.01, \*\*\**p* < 0.001, \*\*\*\**p* < 0.0001.

TSC-deficient cells typically display elevated mTORC1 signaling toward its non-lysosomal substrates, such as S6K1, and paradoxically decreased signaling toward its lysosomal substrates, such as TFEB ^43–45^ (Figures 1C and 2B). Therefore, we reasoned that the mTORC1-dependent effects on TSC1 protein stability may, in turn, feed back into mTORC1 signaling toward its other substrates. Indeed, cells expressing TSC1^2A^—that is less stable than wild-type TSC1—displayed strongly decreased TFEB phosphorylation, compared to TSC1^WT^-expressing cells (Figures 4E-F). Conversely, S6K1 phosphorylation was comparable between TSC1^2A^- and TSC1^WT^-expressing cells (Figures 4E,G), suggesting a substrate-selective mode of mTORC1 regulation downstream of TSC1 phosphorylation. As expected, the downregulation of lysosomal mTORC1 signaling was accompanied by elevated nuclear relocalization of TFE3—another MiT/TFE family transcription factor (Figures 4H-I). Finally, in line with the role of TFEB/TFE3 in promoting the transcription of genes implicated in lysosome biogenesis ^46^, TSC1^2A^-expressing cells showed an overall increase in lysosome abundance, as assessed by LysoTracker staining (Figure 4J). In summary, these data show that mTORC1-mediated phosphorylation of TSC1 stabilizes the protein by preventing its proteasomal degradation and, in turn, TSC1 phosphorylation selectively regulates the lysosomal branch of mTORC1 signaling, thus establishing a lysosomal autoregulatory circuit to fine-tune its activity.

## Discussion

For more than two decades, the mTOR signaling pathway has been known to operate in a linear fashion in which the TSC complex regulates the nucleotide-binding status of RHEB to control the catalytic activity of mTORC1 ^9,10,14^. Here, we demonstrate that mTORC1 also functions immediately upstream of the TSC complex, thus forming a minimal TSC-mTORC1-TSC feedback loop (Figure 4K). The importance of such a short signaling circuit lies in the fact that any dysregulation upstream of the TSC complex, such as alterations in the PI3K/AKT or RAS/MAPK pathways, can potentially be overridden by phosphorylation events on TSC1 that directly modulate lysosomal mTORC1 activity.

Although TSC1 can be phosphorylated by multiple kinases, including PLK1, CDK1, and IKKβ ^47–49^, mTORC1 itself has not been previously described to act directly on the TSC. Here, using different cellular models of mTORC1 hyperactivation, we initially observed an upshift in TSC1 mobility on SDS-PAGE, which we found is due to an increase in its phosphorylation. After confirming that mTORC1 is the upstream kinase responsible for mediating this effect, we identified two mTORC1-dependent phosphosites on TSC1, T1047 and S1080, thereby establishing TSC1 as a novel direct mTORC1 substrate.

How is TSC1 phosphorylation regulated? Under nutrient-replete conditions, a fraction of mTORC1 accumulates on the lysosomal surface, whereas the TSC complex resides mainly in the cytoplasm. Upon nutrient deprivation, this distribution is reversed, with the lysosomal mTORC1 puncta disappearing, and a fraction of TSC relocalizing to lysosomes ^17^. Importantly, the localization of the two complexes on the lysosomal surface is—apparently—not mutually exclusive. First, under basal culture conditions, active mTORC1 dynamically cycles between lysosomes and the cytoplasm on a continuous basis ^50^. Moreover, a fraction of TSC complexes is found associated with lysosomes even in cells grown in nutrient-replete media ^14,17,51^. These previous findings support a model in which active mTORC1 and a fraction of the TSC complex can coexist on the lysosomal surface ^52–54^.

Our data show that disruption of the LAMTOR-Rag supercomplex markedly reduced TSC1 phosphorylation by mTORC1, whereas phosphorylation of S6K1—a non-lysosomal mTORC1 substrate—was largely unaffected, in line with recent work ^33^. Therefore, TSC1 phosphorylation is regulated in a manner similar to that of other lysosomal mTORC1 targets, like TFEB and TFE3. Notably, however, TSC1 phosphorylation is also regulated downstream of growth factor signaling (in addition to AA and glucose availability), is boosted by RHEB overexpression, and is diminished by rapamycin treatment, therefore resembling the regulation of most canonical mTORC1 substrates. Therefore, our data reveal TSC1 as a unique mTORC1 substrate whose regulation combines features of both canonical and non-canonical targets.

Previous studies established that TSC2 stability depends on its interaction with TSC1, which acts as an Hsp90 co-chaperone to prevent HERC1-mediated ubiquitination and degradation ^11,12^. Additional E3 ligases, including members of the TRIM (tripartite motif) family, target both TSC1 and TSC2 for proteasomal degradation ^55^, while sustained AKT activation disrupts the TSC complex and promotes proteasome-dependent TSC2 turnover ^56^. Less is known about the mechanisms controlling TSC1 stability. Here, we report that a phospho-dead TSC1^2A^ mutant exhibits markedly enhanced protein turnover, compared with wild-type TSC1, driven by proteasome-dependent degradation. Notably, this effect is independent of TSC2 binding, as wild-type and the phospho-dead TSC1 mutant are similarly capable of interacting with TSC2. Accordingly, both mTORC1-dependent phosphorylation sites reside at the C-terminal region of TSC1, not overlapping with its coiled-coil domain or with regions required for its association with TSC2 or TBC1D7 ^5,6^. Together, these findings indicate that mTORC1-mediated phosphorylation stabilizes TSC1, independently of TSC complex formation, revealing a protective mechanism by which mTORC1 safeguards its own negative regulator.

Expression of the phospho-dead TSC1^2A^ mutant led to reduced TFEB phosphorylation and increased TFE3 nuclear localization, compared to cells expressing wild-type TSC1, while S6K1 phosphorylation was not significantly different by TSC1 phosphorylation. Such opposing effects on lysosomal versus non-lysosomal substrates have been described previously, wherein TSC loss-of-function models display decreased phosphorylation and enhanced nuclear occupancy of TFEB/TFE3, with a concomitant increase in S6K phosphorylation ^43,44,57^. The inverse pattern is observed in RHEB KO cells that present largely diminished mTORC1 activity (shown as strongly decreased S6K phosphorylation), but elevated TFEB/TFE3 phosphorylation ^50^. In sum, the data presented in our study add to a growing body of work supporting the substrate-selective regulation of mTORC1 downstream of the TSC–RHEB signaling axis.

Collectively, our findings reveal a TSC1-mTORC1-TSC1 feedback inhibition loop that operates locally at lysosomes to coordinate compartmentalized mTORC1 signaling. According to this model, hyperactivation of mTORC1 on lysosomes would cause enhanced phosphorylation and stabilization of TSC1, which, in turn, would suppress lysosomal mTORC1 activity to restore a steady-state transcriptional program downstream of TFEB/TFE3 activation. Such a regulatory signaling circuit enables cells to homeostatically fine-tune anabolic and catabolic cellular processes in response to fluctuations in nutrient and growth factor signals that integrate on lysosomal mTORC1.

## Methods

### Cell culture

All cell lines were grown at 37 °C, 5% CO_2_, cultured in high-glucose Dulbecco’s Modified Eagle Medium (DMEM) (#41965039, Gibco), supplemented with 10% fetal bovine serum (FBS) (#F7524, Sigma; #P30-3306, PAN-Biotech; #FBS.HP.0500, Bio&SELL) and 1x Penicillin-Streptomycin (#15140122, Gibco; #P4333-100ML, Sigma).

Human female embryonic kidney HEK293FT cells were purchased from Invitrogen (#R70007, Invitrogen; RRID: CVCL_6911), human osteosarcoma U2OS cells (#HTB-96, ATCC; RRID: CVCL_0042) were generously provided by Nils-Göran Larsson (MPI-AGE), and control MEFs were generously provided by Kun-Liang Guan. The identity of the HEK293FT cells was validated by the Multiplex human Cell Line Authentication test (Multiplexion GmbH), which uses a single-nucleotide polymorphism (SNP) typing approach, and was performed as described at www.multiplexion.de. All cell lines were regularly tested for *Mycoplasma* contamination, using a PCR-based approach and were confirmed to be *Mycoplasma*-free.

### Cell culture treatments

For amino acid (AA) starvation experiments, custom-made starvation media were formulated according to the Gibco recipe for high-glucose DMEM, omitting all AAs. The media were filtered through a 0.22 μm filter device and tested for proper pH (pH 7.4) and osmolality before use. For the respective treatments under AA-replete conditions, commercially available high-glucose DMEM media were used. All treatment media were supplemented with 10% dialyzed FBS (dFBS) and 1x Penicillin-Streptomycin. For this purpose, FBS was dialyzed against 1x PBS through a 3,500 MWCO (molecular weight cut-off) dialysis tubing. For AA starvation, culture media were replaced with starvation media for 1 hour. For AA add-back experiments, cells were first starved as described above, and then starvation media were replaced with complete media supplemented with 10% dFBS and 1x Penicillin-Streptomycin for 30 minutes. For glucose starvation, cells were cultured for 16 hours in DMEM without Glucose (#11966025, Gibco) supplemented with 10% dFBS and 1x Penicillin-Streptomycin. For glucose add-back, cells were first starved for glucose as described above, and media were replaced by high-glucose DMEM supplemented with 10% dFBS and 1x Penicillin-Streptomycin for 2 hours. For growth factor starvation, cells were cultured for 16 hours in high-glucose DMEM supplemented with 1x Penicillin-Streptomycin, without FBS. For growth factor add-back, FBS was added drop-wise to the cells at 10% final concentration for 10 minutes. For Bafilomycin A1 treatment (#BML-CM110-0100, Enzo), the drug was added to the media to a final concentration of 100 nM for 8 hours. Torin1 (#4247, Tocris Bioscience) was added to the media at a final concentration of 250 nM for 1 hour. For proteasome inhibition, MG132 (#M7449, Sigma) was added to the media at a final concentration of 10 μM for 8 hours. To block protein synthesis, cycloheximide (#239763, Sigma) was added to the media at a final concentration of 100 μM as indicated in the figure legends.

### Antibodies

The custom-made, rabbit polyclonal phospho-specific antibody recognizing TSC1 when phosphorylated at S1080 (phospho-TSC1^S1080^) was produced by immunizing animals with a synthetic KLH (Keyhole Limpet Hemocyanin)-conjugated phospho-peptide corresponding to residues around S1080 of human TSC1: IPTTVG(pS)LPSSKS. The peptide sequence is 100% identical to the respective mouse TSC1 protein sequence. Antibody generation (peptide synthesis, immunization, and affinity purification of rabbit anti-sera) was outsourced to Davids Biotechnologie GmbH (Regensburg, Germany). A list of all primary antibodies used in this study is found in Table S2.

### mRNA isolation and cDNA synthesis

Total mRNA was isolated from cells using a standard TRIzol/chloroform-based method (#15596018, Invitrogen) according to the manufacturer’s instructions. For cDNA synthesis, mRNA was transcribed to cDNA using the RevertAid H Minus Reverse Transcriptase kit (#EP0451, Thermo Scientific) according to the manufacturer’s instructions.

### Plasmids and Molecular Cloning

The pITR-TTP2-bsd (for TSC1 re-expression) and pITR-TTP2-puro (for TSC2 re-expression) vectors were described previously ^50^. Full-length human TSC1 was subcloned from the corrected pRK7-FLAG-TSC1 plasmid (described in ^17^) into the SfiI and NotI restriction sites of pITR-TTP2-bsd. The TSC1^T1047A^, and TSC1^S1080A^ mutants were generated by site-directed mutagenesis using appropriate DNA oligos, and cloned into the BstEII and NotI restriction sites of the pITR-TTP2-TSC1-bsd vector. A vector expressing human TSC1 spanning amino acids 989-1163 was generated by PCR amplification of the respective region from pITR-TTP2-TSC1-bsd, and cloning into the NcoI and NotI restriction sites of a pETM11 vector. The respective pETM11 vector expressing His_6_-4E-BP1 was described previously ^35^. The TSC2^N1643K^ GAP-inactive mutant (GAP^mut^) was generated by site-directed mutagenesis using appropriate DNA oligos, and cloned into the Bsu36I and EcoRV restriction sites of pcDNA3-FLAG-TSC2 (Addgene plasmid #14129). Full-length human TSC2^WT^ and TSC2^N1643K^ were subcloned from the pcDNA3-FLAG-TSC2 expression vectors into the SfiI and ClaI restriction sites of pITR-TTP2-puro.The vector expressing TSC2 lacking the 424 N-terminal amino acids (TSC2^425–1784^; ΔT1BD) was described in ^17^; and the pcDNA3-FLAG-RHEB^WT^ and RHEB^S16H^ expression vectors were described in ^25^. The pRK5-HA-RAPTOR (plasmid #8513) and pSpCas9(BB)-2A-Puro (pX459) V2.0 vectors (plasmid #62988) were purchased from Addgene. All restriction enzymes were purchased from Thermo Scientific. The integrity of all constructs was verified by sequencing. All oligonucleotides used in this study are listed in Table S3.

### Plasmid DNA transfections

Plasmid DNA transfections in HEK293FT cells were performed using X-tremeGENE HP transfection reagent (#06366236001, Roche) in a 3:1 DNA/transfection reagent ratio according to the manufacturer’s instructions. For experiments using RHEB expression vectors, the Effectene transfection reagent (#301425, QIAGEN) was used according to the manufacturer’s instructions. The respective pcDNA3-FLAG, pcDNA3-HA, or pITR-TTP2 empty vectors were used as controls for transfection experiments.

### Generation of knockout cell lines

The HEK293FT TSC1 KO cells were described previously ^58^. The TSC2 KO HEK293FT cell lines were generated using the pX459-based CRISPR/Cas9 method, as described elsewhere ^59^. The sgRNA expression vector was generated by cloning appropriate DNA oligonucleotides (Table S3) into the BbsI restriction sites of pX459 (#62988, Addgene). An empty pX459 vector was used to generate matching control cell lines. In brief, transfected cells were selected with 3 μg/mL puromycin (#A1113803, Gibco) 36-40 hours post transfection. Single-cell clones were generated by FACS-sorting into 96-well plates, and knockout clones were validated by immunoblotting and functional assays.

### Generation of stable cell lines

The polyclonal reconstituted HEK293FT TSC2 KO cell lines stably expressing TSC2^WT^ and TSC2^N1643K^ were generated using the doxycycline-inducible, Sleeping Beauty-based pITR-TTP2 transposon system ^60,61^. In brief, TSC2 KO cells were co-transfected with pITR-TSC2^WT^ or pITR-TSC2^N1643K^ and the transposase-expressing pCMV-Trp vector in a 10:1 ratio. Forty hours post-transfection, cells were selected with 3 μg/mL puromycin (#A1113803, Gibco). The polyclonal cell lines were subsequently maintained in media containing the selection agent. Doxycycline-induced expression from the integrated plasmid was tested by treating the cells overnight with 1 µg/mL doxycycline (#D9891, Sigma). For experiments, all cell lines were used without doxycycline induction to allow for low-level, leaky TSC2 expression.

### Gene silencing experiments

Transient knockdown of *RRAGA, RRAGB, RRAGC, RRAGD,* and *LAMTOR1* was performed using siGENOME (pool of 4) gene-specific siRNAs (Horizon Discoveries). An siRNA duplex targeting the *R. reniformis* luciferase gene (RLuc) (#P-002070-01-50, Horizon Discoveries) was used as control. Transfections were performed using 20 nM siRNA and the Lipofectamine RNAiMAX transfection reagent (#13778075, Invitrogen), according to the manufacturer’s instructions. Cells were collected 72 hours post-transfection and knockdown efficiency was verified by immunoblotting.

### Cell lysis and immunoblotting

For standard SDS-PAGE and immunoblotting experiments, cells from one well of a 12-well plate were treated as indicated in the figures and lysed in 120-200 μL of ice-cold Triton lysis buffer (50 mM Tris pH 7.5, 1% Triton X-100, 150 mM NaCl, 50 mM NaF, 2 mM Na-vanadate, 0.011 gr/mL beta-glycerophosphate), supplemented with 1x PhosSTOP phosphatase inhibitors (#04906837001, Roche) and 1x cOmplete protease inhibitors (#11697498001, Roche), for 10 minutes on ice. Lysates were clarified by centrifugation (14000 rcf, 10 min, 4 °C) and supernatants transferred to a new tube. Protein concentration was determined using a Protein Assay Dye Reagent (#5000006, Bio-Rad). Normalized samples were boiled in 1x SDS sample buffer for 5 min at 95 °C (6x SDS sample buffer: 350 mM Tris-HCl pH 6.8, 30% glycerol, 600 mM DTT, 12.8% SDS, 0.12% bromophenol blue).

Protein samples were subjected to electrophoretic separation on SDS-PAGE and analyzed by standard Western blotting techniques. For achieving maximal electrophoretic resolution of TSC1, lysates were run on a 6% polyacrylamide gel until the 100 kDa protein marker reached the bottom of the gel. In brief, proteins were transferred to nitrocellulose membranes (#10600002 or #10600001, Amersham) and stained with 0.2% Ponceau solution (#33427-01, Serva) to confirm equal loading. Membranes were blocked with 5% skim milk powder (#42590, Serva) in TBS-T [1x TBS, 0.1% Tween-20 (#A1389, AppliChem)] for 1 hour at room temperature, washed three times for 5 min with TBS-T and then incubated with primary antibodies in TBS-T with 5% bovine serum albumin (BSA; #10735086001, Roche or #8076, Carl Roth) overnight at 4°C. The next day, membranes were washed three times for 5 min with TBS-T and incubated with the appropriate HRP-conjugated secondary antibodies (1:10000 in 5% milk in TBS-T) for 1 hour at room temperature. Signals were detected by enhanced chemiluminescence (ECL), using ECL Western Blotting Substrate (#W1015, Promega); or SuperSignal West Femto Substrate (#34095, Thermo Scientific) for weaker signals. Immunoblot images were captured on films (#28906835, GE Healthcare; #4741019289, Fujifilm). Blots were scanned and then quantified using GelAnalyzer 19.1. A list of all primary and secondary antibodies used in this study is provided in Table S2.

### Lambda-phosphatase treatment assays

Cells were lysed in 200 μL ice-cold Triton lysis buffer (50 mM Tris pH 7.5, 1% Triton X-100, 150 mM NaCl) supplemented with 1x EDTA-free cOmplete protease inhibitors (#11873580001, Roche), as described above. Lysates were cleared by centrifugation (14000 rcf, 10 min, 4 °C) and 100 units of λ-phosphatase (#P0753, New England Biolabs) was added to the supernatants, followed by incubation at 30 °C for 30 min. SDS sample buffer at 1x final concentration was added to the reactions, followed by boiling for 5 min at 95 °C. Protein lysates were analyzed by immunoblotting as described above.

### Co-immunoprecipitation (co-IP)

For co-immunoprecipitation experiments, two wells of a near-confluent 6-well plate were lysed in 0.3 mL CHAPS IP buffer each (50 mM Tris pH 7.5, 0.3% CHAPS, 150 mM NaCl, 50 mM NaF, 2 mM Na-vanadate, 0.011 gr/mL beta-glycerophosphate) supplemented with 1x PhosSTOP phosphatase inhibitors (#04906837001, Roche) and 1x cOmplete protease inhibitors (#11697498001, Roche) for 10 minutes on ice. Samples were clarified by centrifugation (14000 rcf, 10 min, 4 °C) and a fraction of the samples was kept aside as input. For anti-HA IPs, 30 μL of pre-washed anti-HA-agarose beads (#A2095, Sigma) were added to the remaining volume of the supernatants and the IP samples were incubated at 4 °C in an overhead rotator for 2 h. For anti-TSC1 IPs, cells were lysed in ice-cold Triton lysis buffer (50 mM Tris pH 7.5, 1% Triton X-100, 150 mM NaCl, 50 mM NaF, 2 mM Na-vanadate, 0.011 gr/mL beta-glycerophosphate), supplemented with 1x PhosSTOP phosphatase inhibitors (#04906837001, Roche) and 1x cOmplete protease inhibitors (#11697498001, Roche), for 10 minutes on ice and a fraction of the samples was kept aside as input. The remaining volume of the supernatants was incubated with 1 µL anti-TSC1 antibody (#6935, Cell Signaling Technology) at 4 °C in an overhead rotator for 3 h, followed by incubation with 30 µL pre-washed Protein A agarose bead slurry (#11134515001, Roche) for an additional hour at 4 °C in an overhead rotator. Beads were then washed three times with CHAPS IP wash buffer (50 mM Tris pH 7.5, 0.3% CHAPS, 150 mM NaCl, 50 mM NaF) or Triton IP wash buffer (50 mM Tris pH 7.5, 1% Triton X-100, 150 mM NaCl, 50 mM NaF) and boiled in 2x SDS loading buffer. Samples were analyzed by SDS-PAGE and the presence of co-immunoprecipitated proteins was detected by immunoblotting using appropriate antibodies.

### Production of recombinant His_6_-tagged 4E-BP1 and His_6_-tagged TSC1^989–1163^ proteins in bacteria

Recombinant His_6_-tagged 4E-BP1 and His_6_-tagged TSC1^989-1163^ proteins were produced by transforming *E. coli* BL21 RP chemically competent bacteria with the respective pETM11-4E-BP1 and pETM11-TSC1^989-1163^ vectors described above, according to standard procedures. In brief, protein expression was induced with IPTG (isopropyl-β-D-thiogalactopyranoside) for 4 hours at 30 °C, and His_6_-tagged proteins were purified using Ni-NTA agarose (#1018244, QIAGEN) and eluted with 250 mM imidazole (#A1073, Applichem).

### mTORC1 *in vitro* kinase (IVK) assays

*In vitro* mTORC1 kinase assays were developed based on previous reports using endogenous mTOR complexes immunopurified from HEK293FT cells ^58,62,63^. In brief, cells of a near-confluent 10 cm dish were lysed in CHAPS IP buffer (50 mM Tris pH 7.5, 0.3% CHAPS, 150 mM NaCl, 50 mM NaF, 2 mM Na-vanadate, 0.011 gr/ml beta-glycerophosphate) supplemented with 1x PhosSTOP phosphatase inhibitors (#04906837001, Roche) and 1x cOmplete protease inhibitors (#11836153001, Roche) for 10 min on ice. Samples were clarified by centrifugation (14000 rcf, 10 min, 4 °C), supernatants were collected and a fraction was kept aside as input material. The remaining supernatants were subjected to immunoprecipitation by incubation with 2 μL anti-mTOR antibody (#2983, Cell Signaling Technology) for 3 hours (4 °C, rotating), followed by incubation with 30 μL of pre-washed Protein A agarose bead slurry (#11134515001, Roche) for an additional hour (4 °C, rotating). Beads were then washed four times with CHAPS IP wash buffer (50 mM Tris pH 7.5, 0.3% CHAPS, 150 mM NaCl, 50 mM NaF) and once with kinase wash buffer (25 mM HEPES pH 7.4, 20 mM KCl). Kinase reactions were prepared by adding 10 μL 3x kinase assay buffer (75 mM HEPES/KOH pH 7.4, 60 mM KCl, 30 mM MgCl_2_) to the beads. Reactions were started by adding 10 μL of kinase assay start buffer (25 mM HEPES/KOH pH 7.4, 140 mM KCl, 10 mM MgCl_2_), supplemented with 500 μM ATP and 35 ng recombinant His_6_-4E-BP1 or 50 ng recombinant His_6_-TSC1^989-1163^ substrates. All reactions were incubated at 30 °C for 30 min, and stopped by the addition of one volume 2x SDS sample buffer and boiling for 5 min at 95 °C. For mTOR inhibition, 250 nM Torin1 was added in the tubes for 10 min at room temperature before starting the IVK reactions. Samples were run in SDS-PAGE, and the mTORC1-mediated phosphorylation on 4E-BP1^T37/46^ and TSC1^S1080^ was detected by immunoblotting with the respective phospho-specific antibodies.

### Immunofluorescence and confocal microscopy for TFE3 localization

For TFE3 staining, cells were seeded on glass coverslips coated with 50 μg/mL fibronectin (#G1393, Sigma) and fixed with 4% paraformaldehyde (PFA) (#28908, Thermo Scientific) in 1x PBS (10 min, room temperature), followed by a permeabilization step with 0.1% Triton X-100 for 10 min. Cells were blocked in 1% BSA in PBS for 45 minutes. Staining with an anti-TFE3 antibody diluted 1:200 in blocking solution was performed overnight at 4 °C. On the next day, coverslips were washed three times with PBS and then stained with highly cross-adsorbed fluorescent secondary antibodies (Donkey anti-Rabbit Alexa Fluor 488, Jackson ImmunoResearch) diluted 1:200 in blocking solution for 1 hour. Nuclei were stained with DAPI (#A1001, VWR) (1:6000 in PBS) for 10 min and coverslips were washed two times for 10 min with PBS before mounting on glass slides with Fluoromount-G (#00-4958-02, Invitrogen). All images were acquired on an SP8 Leica confocal microscope (TCS SP8 X, Leica Microsystems) using a 40x oil objective lens. Image acquisition was performed using the LAS X software (Leica Microsystems). Images from single channels are shown in grayscale, whereas in merged images Alexa Fluor 488 is shown in green and DAPI in blue. Brightness and contrast were adjusted for visualization purposes using Fiji (https://imagej.net/software/fiji/downloads) ^64^. Alterations were applied to the entire image, keeping the parameters identical between all images of the same channel in the panel.

For the assessment of TFE3 nuclear localization, signal intensity was measured using regions-of-interest (ROIs) corresponding to the nuclei of 95 individual TSC1^WT^-expressing cells from 5 independent representative images; and the nuclei of 108 individual TSC1^2A^-expressing cells from 4 independent representative images. Integrated density was calculated using the Fiji software ^64^, representing the sum of the values of all pixels in the given ROI.

### LysoTracker staining and quantification of LysoTracker intensity

For LysoTracker staining experiments, cells were seeded on fibronectin-coated coverslips and grown until they reached 80-90% confluency. Lysosomes were stained by the addition of 100 nM LysoTracker Red DND-99 (#L7528, Invitrogen) in the media for 1 hour. Cells were then fixed with 4% PFA in PBS for 10 min at room temperature, washed and permeabilized with PBT solution (1x PBS, 0.1% Tween-20), and nuclei stained with DAPI (1:2000 in PBT) for 10 min. Coverslips were mounted on slides using Fluoromount-G (#00-4958-02, Invitrogen). All images were captured on an SP8 Leica confocal microscope (TCS SP8 X, Leica Microsystems) using a 40x oil objective lens. Image acquisition was performed using the LAS X software (Leica Microsystems).

Signal intensity was calculated using the Fiji software ^64^. Regions-of-interest (ROIs) were determined for 96 cells from 8 independent representative images per condition and integrated density was calculated, representing the sum of the values of all pixels in the given ROI.

### Phosphosite identification on TSC1 by mass-spectrometry

#### Sample preparation

For the identification of putative mTOR-dependent phosphosites on TSC1, anti-TSC1 IPs were performed as described above using HEK293FT cells (one full 6-well plate per condition) transiently expressing FLAG-tagged RHEB^S16H^ with or without Torin1 treatment (250 nM, 1h) before lysis. After washing the beads in IP wash buffer, immunoprecipitated proteins were washed three times in 50 mM Tris (pH 7.5) and eluted in 100 μL elution buffer [5 ng/μL Trypsin, 50 mM Tris pH7.5, 1 mM TCEP (Tris(2-carboxyethyl)phosphine), 5 mM CAA (chloroacetamide)] for 1 hour at room temperature. After elution, supernatants were transferred to 0.5 mL tubes and incubated at 37 °C overnight to ensure complete tryptic digestion. Digestion was stopped by adding 50% formic acid (FA) to the reaction at a final concentration of 1%. Samples were centrifuged at 20,000 x g for 10 min at room temperature and supernatants were collected. C-18-SD StageTips were washed and equilibrated sequentially with 200 μL methanol, 200 μL 40% ACN (acetonitrile)/0.1% FA and 200 μL 0.1% FA by centrifugation, each step for 1 min at room temperature. Samples were diluted with 0.1% FA, loaded on StageTips and centrifuged for 1-2 min at room temperature. StageTips were then washed twice with 200 μL 0.1% FA. Tryptic peptides were eluted from StageTips with 100 μL 40% acetonitrile (ACN)/0.1% FA by centrifugation (300 x g, 4 min, RT). Eluates were dried in a Speed-Vac at 45 °C for 1 hour and resuspended in 20 μL 0.1% FA. Peptides were stored at -20°C until LC-MS/MS analysis.

### LC-MS/MS analysis

Peptides were separated on a 25 cm, 75 μm internal diameter PicoFrit analytical column (New Objectives) packed with 1.9 μm ReproSil-Pur 120 C18-AQ media (#r119.aq., Dr. Maisch) using an EASY-nLC 1200 (Thermo Fisher Scientific). The column was maintained at 50°C. Buffer A and B were 0.1% formic acid in water and 0.1% formic acid in 80% acetonitrile. Peptides were separated on a segmented gradient from 6% or 3% to 31% buffer B for 45 min at 200 nL/min. Eluting peptides were analyzed on a QExactive HF mass spectrometer (Thermo Fisher Scientific). Peptide precursor m/z measurements were carried out at 60000 resolutions in the 300 to 1800 m/z range. The most intense precursors with charge state from 2 to 7 only were selected for HCD fragmentation using 25% normalized collision energy. The m/z values of the peptide fragments were measured at a resolution of 30000 using an AGC target of 2e5 and, 80 ms maximum injection time. Upon fragmentation, precursors were put on a dynamic exclusion list for 45 sec.

### Protein identification and quantification

The raw data were analyzed with MaxQuant version 1.6.1.0 ^65^. Peptide fragmentation spectra were searched against the reviewed sequences for human or the sequence for TSC1 only. Methionine oxidation, protein N-terminal acetylation, and Phospho (STY) were set as variable modifications; cysteine carbamidomethylation was set as fixed modification. The digestion parameters were set to “specific” and “Trypsin/P”; the minimum number of peptides and razor peptides for protein identification was 1; the minimum number of unique peptides was 0. Protein identification was performed at a peptide spectrum matches and protein false discovery rate of 0.01. The “second peptide” option was on. Successful identifications were transferred between the different raw files using the “Match between runs” option.

To select putative mTORC1 phosphosites on TSC1 for follow-up experiments, the ratio of phospho-peptide intensity to total TSC1 peptide intensity was calculated for each site, and the following filtering criteria were applied: (i) is the site identified as phosphorylated is the ‘RHEB^S16H^’ sample, and (ii) is the phosphorylation robustly decreased in the ‘RHEB^S16H^ + Torin1’ sample? Sites that fulfill both selection criteria were used for downstream analyses.

### Statistical analysis

Statistical analysis and presentation of quantification data was performed using GraphPad Prism (version 10). All relevant information on the statistical details of experiments is provided in the figure legends. Information on quantifications for each method is also provided in the respective Methods section. Data in all graphs are shown as mean ± SD. For graphs with only two conditions shown (Fig. 4I, 4J), significance was calculated using Student’s t-test (unpaired, two-tailed). For all other graphs, significance for the indicated pairwise comparisons was calculated using one-way ANOVA with multiple comparisons test. Sample sizes (n) and significance values are indicated in figure legends (* *p* < 0.05, ** *p* < 0.01, *** *p* < 0.001, **** *p* < 0.0001).

All findings were reproducible over multiple independent experiments, within a reasonable degree of variability between replicates. For most experiments, at least three independent replicates were performed. The sample size for microscopy experiments (number of individual cells used for quantifications) is provided in the respective figure legends. No statistical method was used to predetermine sample size, which was determined in accordance with standard practices in the field. No data were excluded from the analyses. The experiments were not randomized, and the investigators were not blinded to allocation during experiments and outcome assessment.

## Acknowledgements

We thank all members of the Demetriades lab for critical discussions; Aishwarya Acharya and Tânia Catarina Medeiros for feedback on the manuscript; the MPI-AGE FACS and Imaging Core facility for flow cytometry and microscopy support; and the MPI-AGE Proteomics Core facility for support with mass spectrometry work. CD is funded by the European Research Council (ERC) under the European Union’s Horizon 2020 research and innovation programme (grant agreement No 757729), and by the Max Planck Society. Parts of this work were supported by the Deutsche Forschungsgemeinschaft (DFG, German Research Foundation) through the Research Unit Grant FOR2722 (DE 3170/1-1; Project No 384170921) to CD. Models in figures created with BioRender.com.

## Author Contributions

Experimental work: A.L.; data analysis: A.L.; project design, conceptualization: A.L., C.D.; supervision: C.D.; funding acquisition: C.D.; figure preparation: A.L., C.D.; manuscript draft: A.L., C.D. Both authors approved the final version of the manuscript and agree on the content and conclusions.

