## Supplementary material for "TSC1 phosphorylation by lysosomal mTORC1 establishes a minimal autoregulatory feedback loop": Figures S1-S4

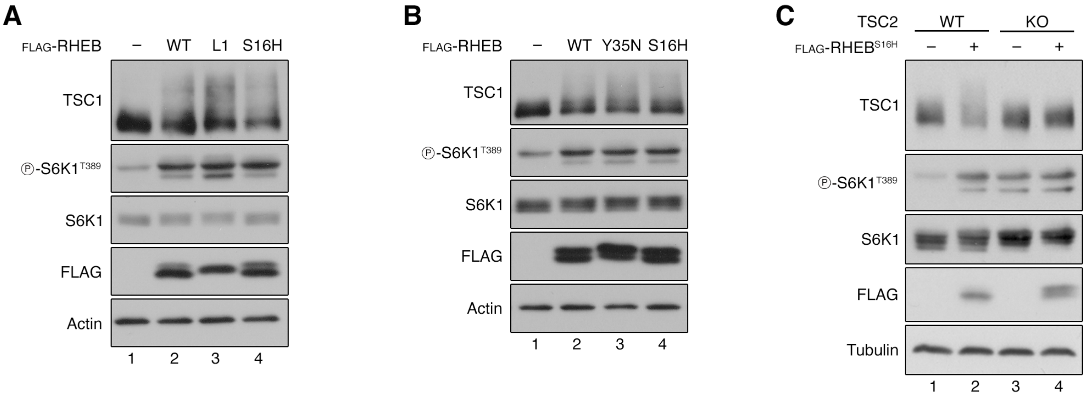


Figure S1

**Figure S1. Exogenous expression of active RHEB or RHEBL1 drives TSC1 phosphorylation in a TSC2-dependent manner. Related to Figure 1.**

**(A)** Immunoblots with lysates from HEK293FT cells transiently expressing FLAG-tagged wild-type RHEB (WT), RHEBL1 (L1), $RHEB^{S16H}$ (S16H), or an empty vector as control (–), probed with the indicated antibodies. n = 3 independent experiments.

**(B)** Immunoblots with lysates from HEK293FT cells transiently expressing FLAG-tagged wild-type RHEB (WT), $RHEB^{Y35N}$ (Y35N), or $RHEB^{S16H}$ (S16H), or an empty vector as control (–), probed with the indicated antibodies. n = 3 independent experiments.

**(C)** Immunoblots with lysates from WT or TSC2 KO HEK293FT cells transiently expressing FLAG-tagged $RHEB^{S16H}$ (+) or an empty vector as control (–), probed with the indicated antibodies. n = 3 independent experiments.

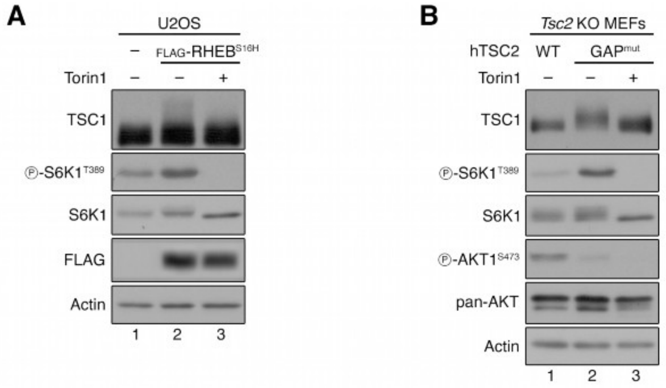


Figure S2

**Figure S2. TSC1 phosphorylation upon mTORC1 activation is not cell-type- or species-specific. Related to Figure 1.**

**(A)** Immunoblots with lysates from U2OS cells transiently expressing FLAG-tagged RHEB$^{S16H}$ or an empty vector, treated with Torin1 (250 nM, 1 h) or DMSO as control, probed with the indicated antibodies. n = 3 independent experiments.

**(B)** Immunoblots with lysates from *Tsc2* KO MEFs transiently expressing WT human TSC2 (hTSC2) or the N1643K GAP-inactive hTSC2 mutant (GAP$^{mut}$), treated with Torin1 (250 nM, 1 h) or DMSO as control, probed with the indicated antibodies. n = 3 independent experiments.

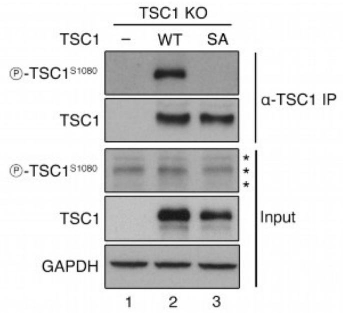


Figure S3

**Figure S3. Specific detection of $TSC1^{S1080}$ phosphorylation in anti-TSC1 IP samples using a custom-made phospho-specific antibody. Related to Figure 2.**

Validation of the custom-made anti-phospho-$TSC1^{S1080}$ antibody using a phospho-dead TSC1 mutant. TSC1 immunoprecipitation from TSC1 KO HEK293FT cells transiently expressing $TSC1^{WT}$ or $TSC1^{S1080A}$ (SA), followed by immunoblotting with the indicated antibodies. Asterisks (*) indicate non-specific bands in blots from whole-cell lysates (Input) using the anti-phospho-$TSC1^{S1080}$ antibody. n = 3 independent experiments.

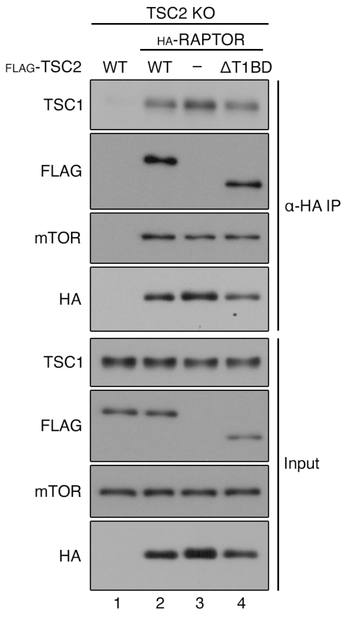


Figure S4

**Figure S4. TSC1-mTORC1 interaction occurs independently of TSC2. Related to Figure 2.**
Co-immunoprecipitation between endogenous TSC1 and exogenously-expressed HA-tagged RAPTOR using lysates from TSC2 KO HEK293FT cells transiently expressing FLAG-tagged TSC2 (WT or a truncate lacking the TSC1 binding domain, ΔT1BD) and HA-tagged RAPTOR. The input and anti-HA IP samples were analyzed by immunoblotting with the indicated antibodies. n = 3 independent experiments.
